# The dentate gyrus controls hippocampal representational drift

**DOI:** 10.1101/2025.09.05.674488

**Authors:** Douglas GoodSmith, Antoine Madar, William H. Carson, Mark E. J. Sheffield

## Abstract

Neural representations evolve across time and experience, a phenomenon termed representational drift.^1–6^ The function and circuit mechanisms of drift remain unknown, including whether it reflects intrinsic biological variability within the population or is actively controlled by circuit inputs.^2,3,7^ Progress has been limited because there is currently no known experimental means to manipulate drift without also altering an animal’s sensory or behavioral experience.^2^ Here we show that drift can be experimentally controlled through targeted manipulation of upstream circuit inputs. We demonstrate this principle in the hippocampus by selectively manipulating dentate gyrus granule cells, which are thought to promote discrimination between similar experiences.^8,9^ Inhibiting granule cells during repeated experiences in familiar virtual environments markedly reduced drift in downstream CA1, revealing that representational drift is not simply an intrinsic property of neural populations, but is actively controlled by upstream circuitry. By experimentally manipulating this circuit-level control of drift, we further discovered that hippocampal drift is concentrated around episode transitions rather than accumulating continuously throughout an experience. These findings establish experimental control over hippocampal drift, revealing a structured, circuit-regulated process that differentiates repeated experiences into distinct episodic representations.

## Main

Place cells in the hippocampus provide a spatial framework that supports episodic memory and spatial navigation.^10^ However, the hippocampus must maintain a balance of stability and flexibility, preserving representations of spatial contexts to support memory retrieval while allowing individual experiences within those contexts to be encoded as distinct memory episodes.^11,12^ Hippocampal population activity has been shown to gradually evolve over repeated exposures to familiar environments, even under highly controlled behavioral conditions.^7,13–15^ This “representational drift” may support memory by allowing the formation of episode-specific memories while preserving a representation of the broader spatial context.^16,17^ However, the circuit mechanisms underlying representational drift remain unclear, including whether drift reflects intrinsic biological variability within the population being recorded or is controlled by inputs from other circuits.^1–3,18–20^ Directly testing the function and dynamics of drift has been challenging because any manipulation that alters an animal’s sensory or behavioral experience also alters hippocampal representations through mechanisms beyond drift itself. Dentate gyrus (DG) granule cells (GCs) are thought to perform pattern separation to support discrimination between similar experiences,^8,9,21^ and are thus well positioned to regulate downstream representational drift between episodes. We therefore performed selective chemogenetic inhibition of GC activity while imaging population activity in dorsal CA1, where hippocampal drift has been well characterized,^7,13,16,17,20,22–24^ to determine whether GC activity controls drift of hippocampal output representations between repeated episodes under constant behavioral and sensory conditions.

### Granule cells drive drift in CA1

To selectively and transiently inhibit GC activity while recording downstream CA1 population activity, we injected a Cre-dependent inhibitory DREADD virus unilaterally into the dorsal DG and GCaMP into dorsal CA1 of Dock10-Cre mice, a mouse line with selective expression of Cre in GCs^25^ (Fig. 1a-b, S1). Imaging from GCs in a separate group of mice confirmed that intraperitoneal injection of the DREADD agonist deschloroclozapine (DCZ)^26^ significantly reduced GC activity (Fig. S2). CA1 activity was recorded while well-trained mice navigated a 2m linear virtual reality (VR) track (Fig. 1c-d, S3) following injection of saline or DCZ.

**Figure 1:**
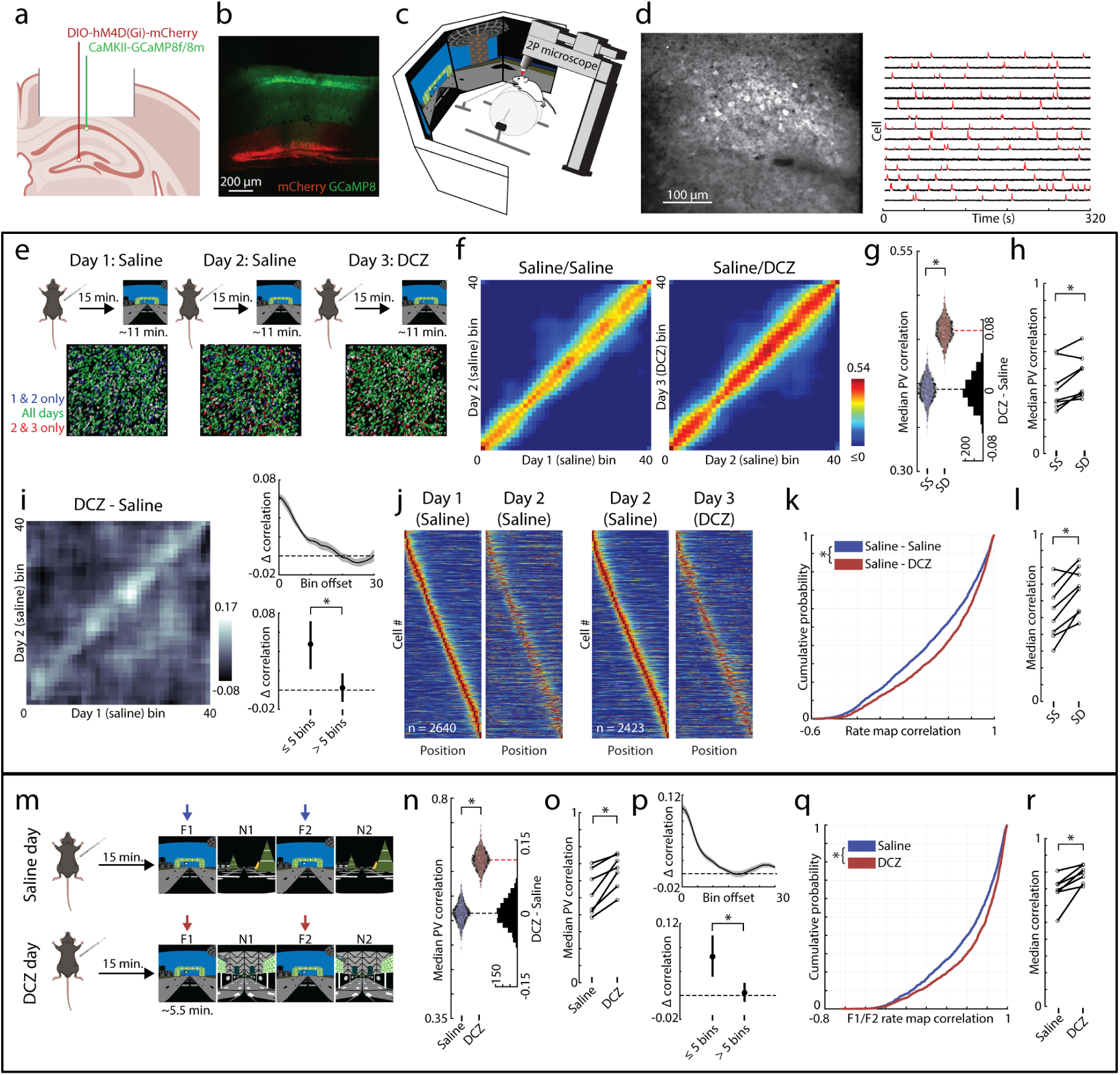
GC inhibition reduces CA1 population drift across episodes in familiar environments. (a-c) Experimental design. A Cre-dependent inhibitory DREADD virus (red) was injected into the DG of Dock10-Cre mice to inhibit GCs, and GCaMP8 was injected into CA1 to image pyramidal cell activity (green). (c) Head-fixed virtual reality imaging setup. (d) Example CA1 field of view (left) and calcium traces with detected transients (red; right). (e) Across-day imaging paradigm (8 mice). Mice received saline injections on days 1 and 2, and DCZ on day 3. The same field of view was tracked, and cells were registered across days. (f) PV correlation matrices across days for saline/saline (control, SS) and saline/DCZ (GC inhibition, SD) conditions. PV correlations at corresponding spatial locations were increased following GC inhibition (SS: n = 4,010 cells; SD: n = 4,086 cells). (g) Bootstrapped difference in median PV correlations (red line) was significantly greater than condition-shuffled values (black histogram of shuffle counts; p < 0.001). (h) GC inhibition increased PV correlations across mice (Wilcoxon signed-rank test p = 0.023). (i)Difference between SD and SS PV correlation matrices (left) and median difference vs spatial offset from main diagonal, showing strongest effects at nearby spatial locations (right; bootstrap permutation test, p = 0.003). (j)Place cell rate maps sorted by peak firing location on the first day show stable spatial representations with detectable drift. (k) CDF plot of place cell rate map correlations across consecutive days. GC inhibition increased correlations (LME, β = 0.17 p = 2.94 x 10^-17^). (l) Increased rate map correlations were consistent across mice (signed-rank test; p = 0.016). (m-r) Same as e-l but for within-day recordings. (m) Mice (n = 7) received saline or DCZ on separate days and were imaged across familiar and novel episodes (F1, N1, F2, N2). (n) GC inhibition increased PV correlations across familiar episodes (saline: n = 6,375 cells; DCZ: n = 7,001 cells; p < 0.001). (o) Increased PV correlations following GC inhibition were consistent across mice (signed-rank p = 0.016). (p) PV correlations were selectively increased at corresponding spatial location (p < 0.001). (q) Place cell correlations between familiar episodes increased following GC inhibition (saline, n = 1,810; DCZ, n = 1,806; LME, β = 0.16, p = 1.02 x 10^-13^). (r) Increased place cell correlations were consistent across mice (signed-rank p = 0.016).

We first aimed to determine whether GC activity regulates the drift of established hippocampal representations across repeated experiences in familiar environments. We considered each re-exposure as a distinct “episode,” and evaluated drift across days (Fig. 1e-l) and across episodes within the same day (Fig. 1m-r). To measure drift across days, we longitudinally tracked CA1 cell populations across 3 days in 8 mice (Fig. 1e) under control (days 1-2; saline/saline) and GC inhibition (days 2-3; saline/DCZ) conditions. Population vector (PV) correlations calculated across all imaged cells (Fig. 1f), a common measure of population-level representational drift,^7,14,24,27^ were significantly increased following GC inhibition (Fig. 1g-h; median [IQR]: saline, 0.39 [0.38-0.41]; DCZ, 0.46 [0.45-0.48]; bootstrap permutation test p < 0.001). The increase in PV correlation was spatially specific, reflecting reduced drift between representations of matched track locations across episodes rather than a global increase in similarity across the entire track (Fig. 1i). We therefore next examined drift at the level of individual place cells by comparing place cell rate map correlation values across days (Fig. 1j-l). Place cell correlations were also significantly increased following GC inhibition (Fig. 1k-l; saline, 0.56 [0.16-0.83]; DCZ, 0.69 [0.31-0.88]; linear mixed effects model (LME), p = 2.94 x 10^-17^). Together, these results establish that GC inhibition reduces across-day drift of established spatial maps.

While representational drift is often measured across days,^7,13,14,24^ its rate is affected by the animal’s experience between episodes.^20,22,24^ In order to control this intervening experience, we also measured drift across episodes within a single continuous experimental session^17,20,27^ in a separate group of mice. Each mouse was imaged on two separate days, receiving either saline or DCZ in random order (different cell populations imaged on each day). On each day, mice alternated between familiar and novel environments (F1, N1, F2, N2; Fig. 1m) to ensure that repeated exposures to the familiar environment (F1 and F2) were treated as distinct episodes rather than a single uninterrupted experience. These within-day recordings also minimized the potential contributions from uncertainty in cell tracking.^7,28^ Consistent with our across-day analysis, GC inhibition significantly increased PV correlations between familiar episodes (Fig. 1n-o; saline, 0.55 [0.54-0.59]; DCZ, 0.68 [0.64-0.69], bootstrap permutation test p < 0.001) selectively at corresponding spatial locations along the track (Fig. 1p). Similarly, GC inhibition increased place cell rate map correlations (Fig. 1q-r; saline, 0.71 [0.36-0.88]; DCZ, 0.79 [0.52-0.92]; LME familiar drug effect p = 1.02 x 10^-13^). These effects were not observed in a separate group of Cre-negative control mice and were not affected by the GCaMP virus used (Fig. S4, S5).

Together, these results demonstrate that GC activity drives drift of established spatial representations across multiple timescales, both across days (separated by ∼24 hours) and within a single day (separated by ∼6 minutes).

### GC-mediated drift is context specific

While GC inhibition reduced drift across episodes in the familiar environment, this effect could reflect a selective regulation of drift of established representations or a general increase in representational stability, including newly formed representations of novel environments. We therefore evaluated drift across episodes (within-day) in novel environments (Fig. 2a-e, S6a-c). Comparing population activity between N1 and N2 provides a measure of the stability of newly formed representations. GC inhibition did not affect novel environment PV correlations (Fig. 2b-c; saline, 0.52 [0.48-0.55]; DCZ, 0.52 [0.49-0.54]; bootstrap permutation test p = 0.97) or place cell rate map correlations (Fig. 2d-e; saline, 0.61 [0.26-0.82]; DCZ, 0.62 [0.25-0.82], LME novel drug effect p = 0.68). The effect of GC inhibition was selective to the familiar environment (Fig. 2f, LME drug x novelty interaction p = 7.59 x 10^-7^), with no effect on remapping between familiar and novel environments (Fig. S6d-e). This result suggests that GCs selectively drive drift of established representations rather than broadly destabilizing newly formed representations.

**Figure 2:**
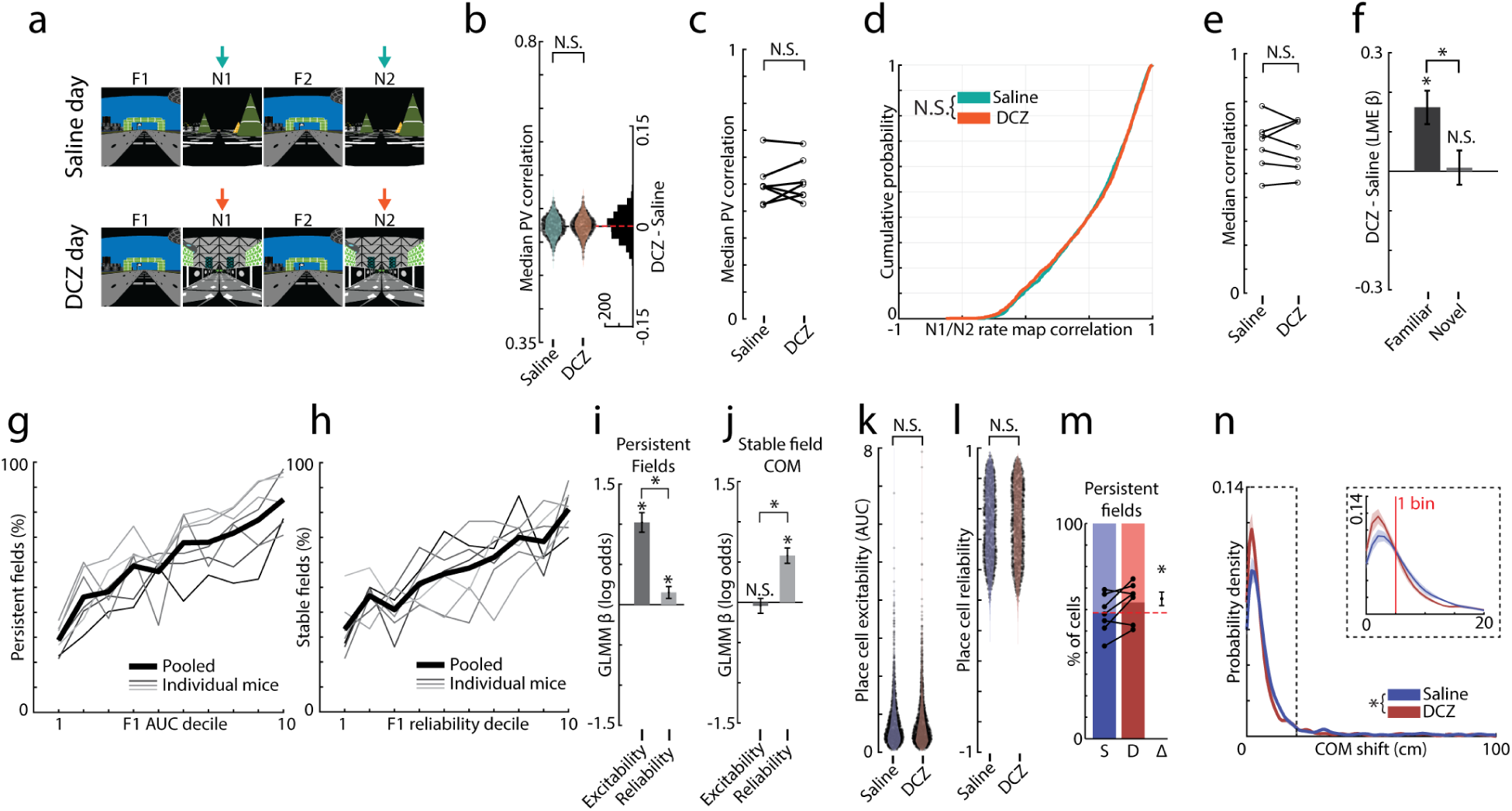
GC-mediated drift is specific to familiar environments and independent of neural correlates of drift. (a-f) Within-day imaging in a novel environment. (a) Spatial stability was assessed between N1 and N2. (b-c) PV correlations were not affected by GC inhibition at the population (b, bootstrap permutation test p = 0.97) or across-mouse (c, signed-rank p = 0.58) levels. (d-e) Place cell rate map correlations were also unchanged (d, LME β = 0.009, p = 0.68; e, signed-rank p = 0.81). (f) The effect of GC inhibition significantly differed between familiar and novel environments (LME novelty x drug interaction β = -0.15, p = 7.59 x 10^-7^). (g-n) Single-cell correlates of drift in the familiar environment. (g)Field persistence increased with F1 place cell excitability (AUC, GLMM β = 1.02, p = 6.97 x 10^-59^). Persistence probability per AUC decile is plotted. Pooled data (thick lines) and individual mice (thin lines), pooled DCZ and saline data (Fig. S7). (h) Spatially stable field probability increased with F1 place cell reliability (GLMM β = 0.58, p = 7.86 x 10^-32^). (i) Place field persistence was more strongly associated with excitability than reliability (GLMM contrast β = 0.87, p = 2.67 x 10^-32^). (j) Among persistent fields, spatial stability was more strongly associated with reliability than excitability (GLMM contrast β = -0.62, p = 4.80 x 10^-17^). (k-l) GC inhibition did not affect place cell excitability (k, LME β = -0.012, p = 0.61) or reliability (l, LME β = -0.006, p = 0.63). (m) GC inhibition increased the probability of field persistence. Individual mice (lines) and GLMM-estimated Δ (β = 0.28, p = 8.23 x 10^-5^). (n) GC inhibition reduced COM displacement across familiar episodes (LME β = -0.36, p = 0.009). Mean and SEM of KDE across mice. Insert: expanded view of small displacements (≤ 20cm) with stable field cutoff (5 cm, red line).

We next asked whether the reduced drift observed in familiar environments could merely reflect changes in single-cell properties previously shown to be correlated with drift.^7,29,30^ Consistent with prior studies,^7,29^ place cell correlations across episodes were positively associated with both excitability (defined as area under the curve (AUC) of the filtered calcium trace) and reliability (defined as the mean pairwise correlations of single lap activity) within the F1 episode (LME, pooled saline and DCZ: AUC p = 1.47 x 10^-42^; reliability p = 1.95 x 10^-211^), with no effect of GC inhibition on either relationship (Fig. S7a; drug x AUC interaction, p = 0.99; drug x reliability interaction, p = 0.23).

Drift can arise either through changes in the active population of place cells or in the spatial locations of place fields across episodes. We therefore decomposed drift into field persistence (the proportion of F1 place cells with a field in F2, regardless of location) and spatial stability (the proportion of persistent fields with the same location in F1 and F2 - center of mass (COM) displacements ≤ 5 cm). Interestingly, field persistence was more strongly associated with excitability in F1 (Fig. 2g, i), while the spatial stability of persistent fields was more strongly associated with reliability in F1 (Fig. 2h, j). However, these relationships were unaffected by GC inhibition (Fig. S7f-i), revealing that GCs do not alter how within-episode excitability and reliability relate to drift across episodes.

Although the relationships between individual cell properties and drift measures were not affected by GC inhibition, GC inhibition could still reduce drift indirectly by increasing the excitability or reliability of active CA1 place cells.^30^ However, GC inhibition had no effect on place cell excitability (Fig. 2k, S7b, d; saline 0.60 [0.36-1.02]; DCZ 0.59 [0.34-0.98]; LME p = 0.61) or reliability (Fig. 2l, S7c, d; saline, 0.50 [0.30-0.70]; DCZ, 0.50 [0.30-0.71]; LME p = 0.63). Instead, GC inhibition independently increased both place field persistence (Fig. 2m; generalized linear mixed effects model (GLMM) p = 8.23 x 10^-5^) and spatial stability (Fig. 2n, S7e; LME p = 0.009) across familiar episodes. These results determine that GCs control drift through changes in the persistence and stability of place fields independently from global modulation of known cellular correlates of drift.

### GCs drive drift at episode transitions

While GC activity selectively drives drift of established representations, it remains unclear when this drift emerges and how it evolves during an experience. Because drift is often evaluated using session-averaged population activity, it is unclear whether drift accumulates gradually throughout an experience or emerges primarily during transitions between episodes.^20,22,27,31^ We therefore examined the temporal dynamics of hippocampal population drift by calculating PV correlations between a reference PV (PVref), created from the F1 episode rate map (first 15 laps) and five-lap rolling average PVs from either F1 or F2 (Fig. 3a, S8a). To evaluate drift within an episode, we first correlated the F1 PVref to lap-wise activity in the same F1 episode (Fig. 3b-d). The difference between saline and DCZ PV correlations (Fig. 3c) was not different from a shuffled distribution (Fig. 3d; p = 0.25) and did not significantly change over the course of the episode (Fig. 3d, S8; p = 0.28), suggesting that GC inhibition did not affect within-episode drift (Fig. S8a-d).

**Figure 3:**
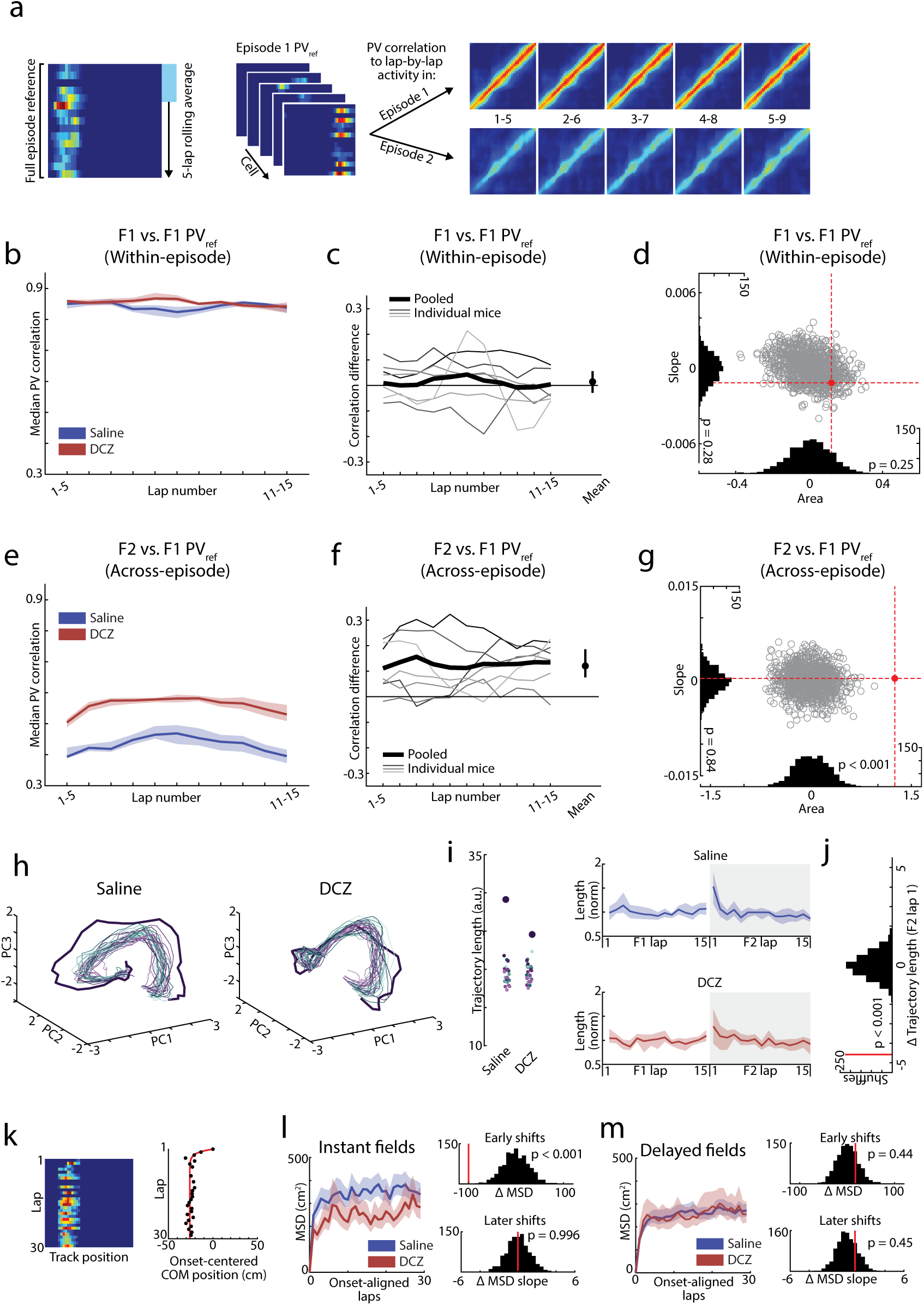
GCs drive drift at familiar episode transitions. (a) Schematic of lap-by-lap PV correlation analysis using an F1 reference PV (PVref). (b-d) Within-episode lap-by-lap PV correlations. (b) PVref correlations with five-lap rolling average PVs from F1. (c) DCZ – saline difference in PV correlations for individual mice (thin lines) and pooled data (thick line). Right, mouse-wise differences did not differ from zero (mean ± CI, signed-rank p = 0.58). (d) Area between saline and DCZ PV curves (x-axis) and the slope of the difference across laps (y-axis). Observed values (red) and 1,000 condition-shuffled values (gray, black histograms). (e-g) Across-episode lap-by-lap PV correlations. (e)F1 PVref correlations with five-lap rolling average PVs from F2. (f)DCZ – saline difference in PV correlations. Right, the mouse-wise mean difference across laps was greater than zero (signed-rank p = 0.016). (g)The area between saline and DCZ PV curves exceeded all shuffled values (p < 0.001), while there was no significant slope in the difference across laps (p = 0.84). (h)PCA trajectories of individual F1(green) and F2 (purple) laps projected into a shared population space. F2 lap 1 is shown in bold. (i) Left, trajectory lengths in PCA space for individual laps. Right, mean ± CI of normalized trajectory length across mice. There was a transient increase in trajectory length on lap 1 of F2. (j) The DCZ – saline difference in F2 lap 1 trajectory exceeded all condition-shuffled values (p < 0.001). (k) Left, example shifting place cell rate map. Right, lap-by-lap COM relative to onset with nonlinear exponential fit (red). (l-m) Place field shifting for instant (l; saline n = 2,487, DCZ = 1,236) and delayed fields (m; saline n = 2,899, DCZ n = 1,135). Left, onset-aligned MSD. MSD increased rapidly during the first laps, reflecting fields shifting early after field onset. This increase was followed by a shallower slope reflecting the extent of ongoing field displacement throughout the episode. Right, DCZ – saline differences in MSD during the first 3 laps (early shifts) were greater than condition shuffled values (p < 0.001), while there was no difference in the slope of MSD values for laps 4+ (later shifts, p = 0.996). Delayed fields were not affected by GC inhibition (m, early shifts p = 0.44, later shifts p = 0.45).

In contrast, GC inhibition significantly increased correlations between F1 PVref and lap-wise activity in F2 (Fig. 3e-g). The difference between saline and DCZ PV correlations was significantly greater than values obtained from a shuffled distribution (Fig. 3g; p < 0.001), suggesting that GC inhibition selectively affected across-episode drift of well-established familiar representations (Fig. S8). Notably, this effect was already apparent from the earliest F2 laps and did not significantly evolve throughout F2 (Fig. 3f-g; p = 0.84). These results establish that GC-mediated drift does not accumulate throughout an episode but instead reflects a discrete change upon re-entry to the familiar environment. Consistent with these results, a PV-based location decoder^32,33^ showed reduced across-episode decoding errors, with no effect on within-episode decoding accuracy, following GC inhibition (Fig. S9).

As GC-mediated drift is concentrated at the start of episodes, we next examined how familiar episode transitions reshape population activity using PCA. PV activity from individual F1 and F2 laps was projected onto principal components derived from the combined population activity across both episodes (Fig. 3h). The first lap of F2 (F2 lap 1) had longer PCA trajectories than all other laps (Fig. 3h-j, S10), with significantly longer trajectories in saline than DCZ conditions (Fig. 3j, S10; permutation test p < 0.001), suggesting that GC activity promotes transient exploration of neural activity states during re-entry to a familiar environment.

We next asked how the population level drift at episode transitions was reflected in the lap-by-lap dynamics of individual place cells. We quantified shifts in place field COM, a measure associated with behavioral timescale synaptic plasticity (BTSP)-mediated place field dynamics (Fig. 3k).^15,34^ Across-day recordings were used to ensure sufficient laps for analysis (saline episode 1 laps: within-day 25 [23 – 32]; across-day 49 [39 – 63]; Fig. S3). GC inhibition reduced place field shifting selectively among place fields that were active from the first lap (instant fields; Fig. 3l), with no effect on fields that emerged later in the episode (delayed fields; Fig. 3m, S11). Notably, the GC-mediated reduction in instant field shifting occurred primarily over the initial laps in the episode (Fig. 3l), suggesting that GCs primarily affect the early evolution of place fields present immediately upon re-entry to the familiar environment.

Together, these results reveal that GC-mediated drift emerges rapidly at episode transitions rather than accumulating gradually throughout an experience. Upon re-entry to a familiar environment, GCs transiently drive CA1 population activity away from established neural activity states, reflected in early shifts of instant place fields. This rapid accumulation of drift at episode transitions may serve to segment distinct episodes and support discrimination between similar experiences.

### Drift supports neural discrimination of episodes

To determine whether GC-mediated drift promotes neural discrimination between repeated experiences, we used a PV template-matching decoding strategy^32,35^ to test whether CA1 population activity could distinguish between episodes occurring in the same spatial context and whether this discrimination was affected by GC inhibition. Although F1 and F2 occurred in the same environment and had substantially overlapping spatial representations (Fig. 1n, q), the decoder was able to reliably distinguish between episodes on saline-injected days, with accuracy approaching the values obtained when decoding between familiar and novel contexts (Decoder accuracy: F1/F2 97.5%, F1/N1 99.6%; Fig. 4a-c, S12). GC inhibition more than tripled the proportion of episode misclassifications (F1/F2 decoder errors: saline 2.5%; DCZ 8.1%; Fig. 4a-c), with the effect of DCZ exceeding all condition-shuffled values (Fig. 4d, permutation test p < 0.001). We further quantified episode separation using a correlation modulation index, defined as the normalized difference between F1 and F2 template correlation values. This metric provides a continuous measure of the degree of episode separation. Consistent with decoder accuracy results, GC inhibition led to a significant decrease in modulation index at the population level (Fig. 4e, permutation test p < 0.001) and across individual mice (Fig. S12).

**Figure 4:**
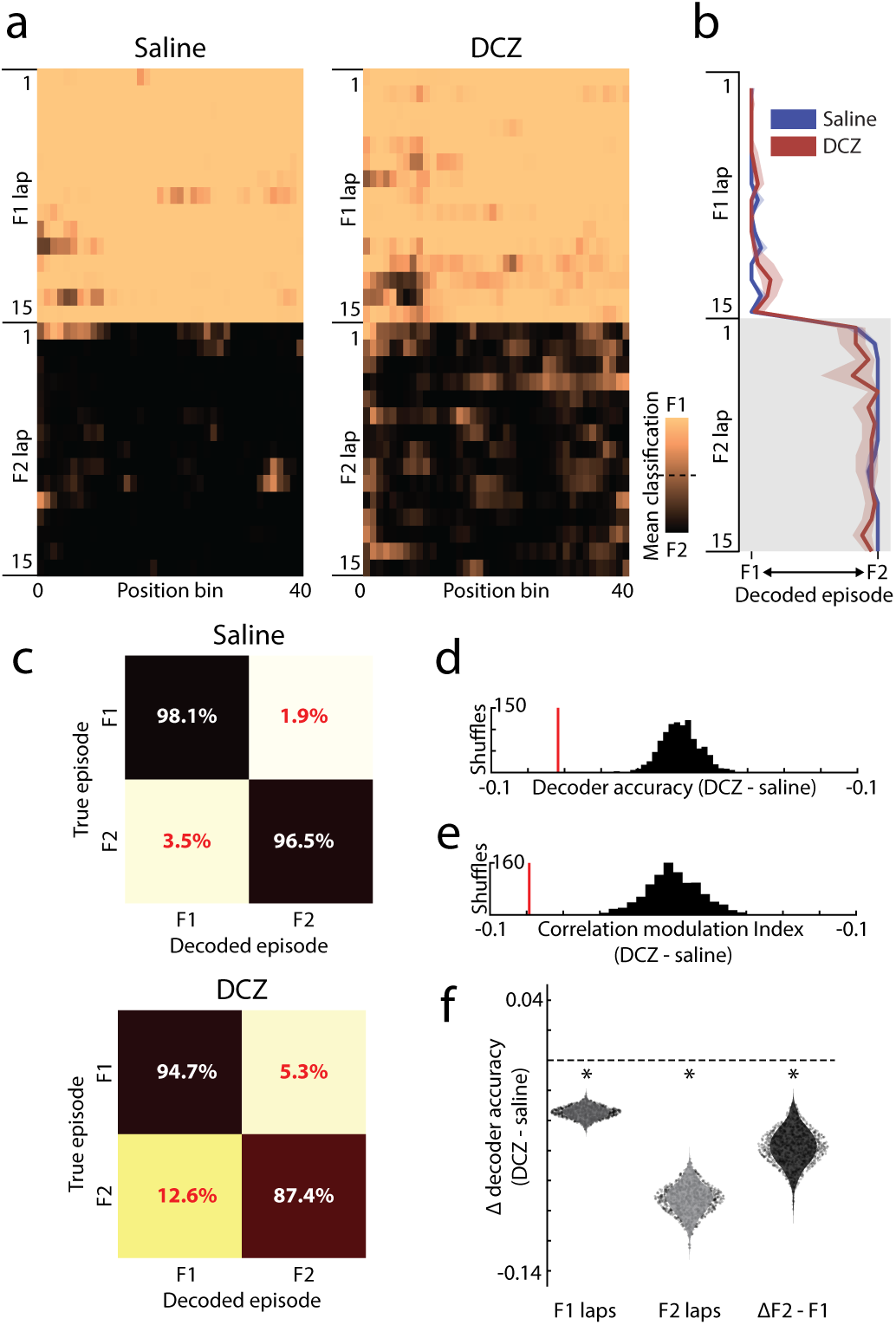
GC-mediated drift supports differentiation between repeated experiences. (a-b) PV-template matching decoding of familiar episodes. (a) Heatmap showing the proportion of decoder iterations classified as F1 or F2 for each lap (row) and spatial bin (column). Lighter colors indicate more F1 classifications, and darker colors indicate more F2 classifications. (b) Decoded episode across laps (median and IQR). (c) Confusion matrices for saline and DCZ conditions. GC inhibition increased episode misclassifications, with errors biased toward classifying F2 episodes as F1. (d) The effect of GC inhibition (DCZ – saline) in decoder accuracy across all laps was significantly greater than all condition-shuffled values (permutation test, p < 0.001). (e) Correlation modulation index was reduced following GC inhibition (permutation test, p < 0.001). (f) GC inhibition reduced mean decoder accuracy in both F1 and F2 episodes, with a larger effect in F2 than F1 (permutation tests relative to condition-shuffled values; F1, p < 0.001; F2, p < 0.001; F2 – F1 interaction, p = 0.006).

GC inhibition significantly reduced decoder accuracy in both F1 and F2 episodes relative to a condition-shuffled distribution (Fig. 4f; permutation tests, p < 0.001). However, F2 episodes were more frequently misclassified as F1 than the reverse (Fig. 4c, f), and the reduction in decoder accuracy was significantly greater for F2 than F1 (Fig. 4f, permutation test p = 0.006). This asymmetry suggests that GC inhibition preferentially constrained the deviation of F2 population activity from the established F1 representation, rather than broadly disrupting decoder performance. Together, these results suggest that GC-mediated drift promotes the formation of distinct episode-specific representations to support discrimination between repeated experiences in the same environment.

## Discussion

In this study, we demonstrate that hippocampal representational drift can be experimentally controlled through manipulation of dentate GCs. GC inhibition selectively reduced drift of established CA1 representations upon re-entry to a familiar environment, while within-episode stability and the stability of novel spatial representations were unaffected. Together, these results identify GCs as a circuit mechanism that drives the divergence of hippocampal representations at episode transitions, allowing repeated experiences to be segmented into distinct episodic representations without disrupting broader contextual representations.

Stable hippocampal representations have long been viewed as essential for reliable memory retrieval, but recent studies have reported considerable drift even in identical environments.^7,13,20,24^ Drift has been suggested to support memory updating or discrimination, but whether drift is an actively regulated process or a consequence of intrinsic biological variability remains unknown.^2,3,18,19,23^ Recent work has shown that drift persists despite stable sensory, behavioral, and experimental conditions,^7^ but manipulating drift without altering these conditions remains difficult as any manipulations that alter an animal’s sensory experience inevitably alter hippocampal representations independently of drift.^7,36^ Consequently, the study of representational drift has largely been confined to observational studies on cellular and behavioral correlates of drift rather than direct tests of the functional role of drift and the circuit mechanisms that regulate it. By altering the magnitude of representational drift independently of the animal’s sensory and behavioral experience through selective hippocampal circuit manipulation, we provide direct evidence that drift can be experimentally controlled and that GC activity drives drift in hippocampal representations. GC manipulation therefore provides an experimental tool for causally testing the computational and mnemonic functions of drift in future studies.

Representational drift is often viewed as the gradual accumulation of small changes in population activity across time.^13,20,24^ Although individual neurons may display more abrupt changes, population-level drift is thought to reflect continual updating rather than discrete representational changes.^20,24,27^ However, drift is often quantified using session-averaged activity that may obscure the temporal dynamics of drift. Our lap-by-lap analyses suggest that GC-mediated drift was largely concentrated around the transitions between episodes, with relatively little effect of GC inhibition on drift occurring during an episode. In addition, in our across-day recordings, inhibiting GCs only during re-exposure to the familiar environment was sufficient to reduce drift, suggesting that GCs primarily regulate the reinstatement of previously established representations. Together, our results establish that GC-mediated drift does not accumulate continuously throughout an experience but is selectively expressed early in an experience as episode-specific representations rapidly emerge to segment similar experiences.

While often treated as a single process, evidence increasingly suggests that representational drift may comprise multiple related but dissociable processes. Previous studies have reported separable effects of experience and elapsed time on drift at different timescales,^20,24^ as well as differences in the characteristics of drift between familiar and novel environments.^37^ We found that distinct components of drift were associated with distinct cellular features, with neuronal excitability^7,30^ more associated with place field persistence and within-episode lap-by-lap reliability^29^ correlated with place field displacement across episodes. GC inhibition reduced lap-by-lap place field shifting during the first laps of an episode, with little effect on place field shifts later in the episode. Within-episode place field shifts have been linked to BTSP, which can occur even during repeated experiences within familiar environments.^15,34^ Whether GCs drive drift upon re-entry to the familiar environment through similar BTSP mechanisms remains to be tested. Together with the context-specific effects of GC inhibition, these results suggest that different forms of drift may be supported by separate circuit mechanisms. The GC-mediated drift we report here may primarily reflect rapid across-episode changes that support segmentation of repeated familiar experiences and most closely resembles the form of drift characterized across exposures to a familiar context in prior studies.^7,13,16,17,20,24^ In contrast, gradual within-episode drift may support memory updating and ongoing learning through distinct mechanisms.^15,20,27,34^

The known circuit functions of GCs provide a candidate mechanism through which GCs could regulate representational drift. Sparse GC activity is thought to support pattern separation to prevent interference between similar experiences.^21,38–41^ Small differences in non-spatial cues and internal motivation between episodes may be reflected in GC activity, where cue,^42^ object,^43^ and reward^44^ related activity has been reported. These changes in GC activity may bias CA3 away from previously established activity patterns, introducing subtle variability to drive network activity away from established attractor basins upon entry to an environment. GC inhibition may reduce the influence of decorrelated GC inputs, allowing CA3 pattern completion to more faithfully reinstate previously established representations and reduce drift in CA1. Although our manipulation primarily targets mature GCs,^45^ the birth and integration of new adult-born GCs could further promote experience-dependent modulation of drift magnitude.^1^ Consistent with this idea, chronic ablation of neurogenesis increases GC place cell correlations across experiences separated by one week.^46^ In contrast, developmental loss of the DG impairs CA1 spatial stability,^47^ possibly reflecting circuit compensation. Our results suggest that the magnitude of representational drift may reflect the balance between pattern separation and pattern completion within the hippocampal circuit, with GC activity promoting the emergence of episode-specific representations.

We propose that GC-mediated pattern separation drives the divergence of established hippocampal representations, supporting segmentation of repeated experiences in the same environment into distinct memory episodes while preserving the broader contextual representations that link them. Notably, representational drift has also been observed across multiple cortical areas,^4–6^ and pattern separation-like computations that decorrelate inputs exist throughout the brain.^48–50^ Our results raise the possibility that controlled representational drift is driven by pattern separation mechanisms in upstream circuit inputs, providing a general strategy through which neural circuits can balance stable representations with the ability to flexibly differentiate between similar experiences.

## Methods

### Subjects

Male and female Dock10-Cre mice^25^ (gift from Tonegawa lab) were used for this study (12-16 weeks old, 20-35g at time of surgery). In total, we recorded from 13 Dock10-Cre mice (7 male and 6 female) and 3 Cre-negative littermates (2 male, 1 female). Mice were group housed on a reverse 12-hour light/dark cycle. All experimental procedures were performed during the animal’s dark cycle. All surgeries and animal procedures complied with National Institutes of Health guidelines and were approved by the Institutional Animal Care and Use Committee at the University of Chicago.

### Surgical procedures

Mice were anesthetized with isoflurane (∼3% induction, ∼1-2% maintenance) and injected with saline (0.5 ml, intraperitoneal) and Meloxicam (1-2 mg/kg, subcutaneous) before being mounted to a stereotaxic surgical station (David Kopf Instruments). A craniotomy was drilled over the right hippocampus (2.0 mm posterior, 1.25 mm lateral of Bregma). Viruses were diluted 1:1 in PBS and injected through a beveled glass micropipette. To target the DG, 150 nl of a Cre-dependent inhibitory DREADD virus (hSyn-DIO-hM4D(Gi)-mCherry; Addgene, #44362) was injected 2.0 mm below the surface of the brain. To target CA1 for imaging, 75nl of GCaMP8f (CAMKIIa-jGCaMP8f; Addgene, #176750) or GCaMP8m (CAMKIIa-jGCaMP8m; Addgene #176751) was injected 1.25 mm below the surface of the brain. Following each injection, the pipette was left in place for 10 minutes before retraction. After both injections, the skull was covered with dental cement (Metabond, Parkell Corporation) and a titanium headplate (Atlas Tool and Die Works) was attached. Following surgery, mice were water restricted and handled daily.

5-8 days after the virus injection surgery, mice underwent a second surgery to implant an imaging window/cannula.^15,51,52^ A small glass circle (2.5 mm diameter, Potomac Photonics), was glued to one side of a stainless-steel cannula (2.77 mm OD, 2.31 mm ID, 1.5 mm length, MicroGroup, Inc.) with UV-curing epoxy (NOA81, Norland). The headplate was removed and a 2.7-2.8 mm craniotomy was made with a trephine drill (Fine Science Tools). The dura was removed and the cortex overlying the hippocampus was aspirated before inserting the cannula. The cannula was secured and a new headplate was attached with Metabond. A 19-mm diameter metal ring was attached to the headplate to house the microscope objective and block out ambient light during imaging. After ∼7 days of recovery, mice began behavior training.

### Behavior and training

Behavior training and imaging was performed in a virtual reality (VR) setup as previously described.^15,51,52^ Mice navigated through 2m long virtual linear environments (created using VIRMEn^53^) displayed on 5 bezel-free monitors. Head fixed mice ran on a Styrofoam or 3D-printed wheel (treadmill). Movement was measured with a rotary encoder (US Digital) and used to update the VR position. Following each traversal (“lap”) of the track, a 4 µl water reward was delivered using a solenoid (NResearch). After a 1.5 second pause, mice were teleported to the start of the track. Licking was detected with a capacitive touch sensor (AT42QT1010, Sparkfun electronics), and movement, reward delivery, and licking data were recorded and synchronized to imaging frames using a Picoscope digital oscilloscope (PICO4824, Pico Technology). Mice were trained daily (∼30 minutes per day) until running 3-5 laps per minute without extended pauses (∼10-14 days). Mice were habituated to IP injections and the imaging objective on the final 3-4 days of training.

### DREADD inhibition

The ligand Deschloroclozapine dihydrochloride (DCZ, MedChemExpress) was used to activate the inhibitory DREADD receptor hM4D(Gi).^54^ Mice were injected with 0.05 – 0.1 ml of a 0.04 mg/ml DCZ solution (0.1 mg/kg, intraperitoneal) 15 minutes before the start of imaging. On separate days, mice were injected with an equal volume (0.05 – 0.1 ml) of saline 15 minutes before the start of imaging. The DCZ dose used in this study (0.1 mg/kg) has been previously used to produce robust DREADD-mediated inhibition in vivo^55–57^ and GC recordings from 2 mice confirmed that DCZ significantly reduced GC activity in this study (Fig. S2).

### Two-photon imaging

Imaging was performed using a laser scanning two-photon microscope (Neurolabware) at 31 Hz with bidirectional scanning through a 16x/0.8 NA/3 mm WD water immersion objective (MRP07220, Nikon). GCaMP8f/8m was excited at 920 nm with a femtosecond-pulsed two photon laser (Insight DS + Dual, Spectra-Physics or Spark lasers Alcor 920-2; 50-100 mW laser power at objective). Emitted fluorescence was collected using GaAsP PMTs (H11706, Hamamatsu). Time-series images were collected through Scanbox (v4.1, Neurolabware).

For across-day experiments (n = 8 mice; Fig. 1e-l), the same FOV was tracked across 3 days by aligning each days imaging plane to a series of reference FOVs collected on day 1. Each day consisted of a single ∼11 minute session in the familiar environment, with saline injected on days 1 and 2 and DCZ on day 3 (15 minutes before imaging). GCaMP8m was used for all 8 mice, and 6,216, 6,034 and 6,045 cells were recorded on days 1-3, respectively.

For within-day experiments (n = 7 mice; Fig. 1m-r), FOVs were selected on each day to maximize cell yield and were not matched across days. Each day consisted of four 10,000 imaging frame (∼5.5-minute) episodes, alternating between a familiar (F1, F2) and novel (N1, N2) environment in the sequence F1-N1-F2-N2. For each mouse, data was analyzed from one saline and one DCZ -injected day (in random order). Days with visible z-drift or failure to run at least 10 laps in all four episodes were excluded, and a replacement day of data was collected in the same condition. The familiar environment was always the same, while a new novel environment was used each day. A total of 6,375 cells on the saline-injected day and 7,001 cells on the DCZ injected day were recorded. Two mice were included in both experiment groups, with the within-day (FNFN) recordings occurring following the across-day recordings. Five mice were recorded using GCaMP8f and two with GCaMP8m. Results did not differ between indicator groups and were therefore pooled for analysis (Fig. S4).

### Perfusion and histology

Mice were anesthetized with isoflurane and perfused with PBS followed by 4% PFA. Brains were stored in PFA overnight and in 30% sucrose for 3-5 days before being sectioned (30-50 µm) on a cryostat (Leica CM 3050S). Sections were mounted with DAPI-Fluoromount (SouthernBiotech) and imaged with an Olympus VS200 Research Slide Scanner (Olympus/Evident, Center Valley, PA) at the Integrated Light Microscopy Core at the University of Chicago (RRID: SCR_019197).

### Image processing and ROI selection

Time-series images were preprocessed using Suite2p.^58,59^ Movement artifacts were removed using rigid and non-rigid transformations and datasets with visible z-drift were excluded. Regions of interest (ROIs) were identified and classified as cells or non-cells using Suite2p with manual curation. ΔF/F traces of cells were baseline corrected and filtered for significant calcium transients as previously described.^15,60^ Briefly, fluorescence values were normalized by the 8^th^ percentile within a 15 second sliding window and median-subtracted. Significant calcium transients were identified with an amplitude and frequency-based analysis with <0.1% false positive rates.^60,61^ Fluorescence values outside of significant events were set to 0. Imaging data were aligned to behavior data and filtered to remove periods of immobility or slow running (< 2 cm/s).

For across-day drift experiments, cells were registered across days using CellReg^28^ with day 1 as the reference. The spatial correlation method was applied to all datasets, and registrations were visually validated. Of all recorded cells, 3,166 could be tracked across all 3 days, and 4,930 were tracked across at least 2 consecutive days. We analyzed cells registered on consecutive days (days 1-2 for saline/saline comparison, n = 4,010; days 2-3 for saline/DCZ comparison, n = 4,086).

## Data analysis

### Activity rate maps and place cell detection

Activity was binned into 40 spatial bins (5 cm) to create lap-by-lap and session averaged activity maps (“rate maps”). Lap-by-lap rate maps were used to identify place field onsets, calculate within-episode reliability, analyze within-episode population dynamics, and to calculate lap-wise PV correlations. Session-averaged maps were used to visualize population place cell activity, calculate spatial information scores, generate session-wide PVs, and calculate rate map correlations. Cells were classified as place cells if they had both a significant spatial information score^62–65^ and a defined onset lap.^15^ Spatial information significance was assessed relative to 100 spatially shuffled maps (random offset between activity and location) for each cell. Cells with observed spatial information scores that exceeded > 95% of shuffled map values were considered significant. To identify place field onsets,^15^ putative fields were defined as contiguous bins exceeding 15% of the difference between the most active bin and the average of the 10 least active bins. Fields separated by a single bin were merged, and putative fields larger than 3 bins (15 cm) with activity on at least five laps were further analyzed. Place field onset was defined as the first lap with activity in the field followed by activity on at least one of the next five laps.

### Place field stability and reliability measures

Within-episode reliability was quantified as the mean of pair-wise Pearson’s correlation coefficients of lap-by-lap activity maps across all laps.^66^ As an additional measure of reliability, we also calculated the correlation between each cell’s average activity during the first and second halves of each episode. Excitability was quantified as the AUC of the filtered, normalized calcium trace, divided by running time (excluding pauses). As a complementary measure, we calculated the proportion of laps that contained at least one significant calcium transient.

As drift between episodes can reflect individual cells gaining or losing place fields, or changes in the spatial locations of fields, we separately quantified these features and their relationship to single-cell excitability and reliability measures. We identified cells with fields in both F1 and F2 and calculated the displacement between place field centers of mass (COMs)^15^ across episodes after circularizing the track. For cells with multiple fields, the minimum displacement across all field pairs was used. We compared cells with persistent fields (detected in both F1 and F2, regardless of location) to cells that had a field in F1 but not F2. Likewise, among persistent fields, we compared spatially stable (COM displacement ≤1 bin) and unstable (COM displacement > 1 bin) fields. For visualization, cells were divided into deciles based on excitability or reliability, and the proportion of persistent vs non-persistent (or stable vs unstable) cells was calculated within each decile.

### Population vector and rate map correlations

Population vectors (PVs) were created for each of the 40 spatial bins using the average activity of all cells, including non-place cells, to capture ensemble-level changes in CA1 spatial representations and avoid biases from place cell identification.^65,67^ Pearson’s correlation coefficients between all pairs of PVs across episodes were calculated to generate PV correlation matrices. Analyses focused on corresponding spatial locations (main diagonal). For across-day recordings, PVs were generated from normalized rate maps (each cell normalized to its peak activity) to account for day-to-day variability in signal amplitude. Spatial selectivity of DCZ-induced PV correlation changes was assessed by subtracting the saline correlation matrix from the DCZ matrix and comparing median correlation differences between matched/nearby locations (≤5 bins) and more distant locations (>5 bins) from the main diagonal.

The effects of GC inhibition on PV correlations were assessed using a bootstrap permutation testing procedure. To control for differences in cell numbers across mice, cells were downsampled with replacement to the minimum number recorded across mice (150 for across-day recordings, 500 for within-day recordings). Differences between median saline and DCZ PV correlations and between condition-shuffled PV correlations (each cell randomly assigned to saline or DCZ) were calculated. This procedure was repeated 1,000 times, with a new bootstrapped cell sample for each iteration. For visualization, (Fig. 1g, n, 2b), bootstrap correlation values are plotted (violin plot) and the shuffle difference distribution (histogram) is centered on the mean saline value. Two-sided permutation test p values were calculated as the proportion of shuffled differences with an absolute value exceeding the mean observed difference value.

To quantify rate map correlations, Pearson’s correlation coefficients were calculated between each cell’s mean rate map across episodes/days, restricted to cells with a place field detected in the first episode/day. The significance of place cell correlation effects was assessed using linear mixed effects (LME) models (see below).

### Mixed Effects models

To account for the nested structure of cells within mice, we used mixed effects models with a random intercept for MouseID for all single-cell analyses unless otherwise stated.^68^ Linear mixed effects (LME) models were used for continuous measures. Within-day place cell correlation analyses included novelty, drug, and their interaction as fixed effects. Across-day analysis included drug as a fixed effect. Correlation coefficients (place cell correlations and reliability) were Fisher z-transformed before analysis. Associations between excitability or reliability and binary measures of place field persistence and stability were assessed using generalized linear mixed-effects models (GLMMs) with a binomial distribution and logit link.^68^ Excitability, reliability, drug, and their interactions with drug were included as fixed effects.

Excitability and reliability were z-scored before modeling. Contrast vectors were used to estimate specific effects and p values, and plots show model estimates and 95% confidence intervals.

### Lap-wise PV analysis

To evaluate lap-by-lap PV dynamics (Fig. 3a-g, S8), a reference PV (PV_ref_) was generated from the average population activity across the first 15 laps of the F1 episode (the minimum number of laps recorded across all mice). Rolling average rate maps were then created using a five-lap moving window in either F1 (within-episode) or F2 (across-episode). Each rolling-average PV was correlated with PV_ref_ at corresponding locations, and the median correlation across bins was used to produce a lap-by-lap PV correlation curve for each condition. To quantify the effects of GC inhibition, the saline PV correlation curve was subtracted from the DCZ curve. The area under the difference curve and slope of the difference across laps were evaluated using permutation testing (see Statistics).

### Place field shifting analysis

Non-linear, lap-by-lap place field shifting was analyzed as previously described.^34^ For each place field, activity outside of the field boundaries used for onset detection was removed to create a field-specific rate map. COM location was interpolated for laps without activity, and any laps following the last defined COM were excluded. Fields with activity on fewer than 30% of all laps or fewer than 30 laps after interpolation were excluded from analysis. COM trajectories were aligned to field onset and truncated at 30 laps. The mean squared displacement (MSD) for each lap was defined as the onset-centered squared displacement across place fields. MSD trajectories were analyzed separately for fields active from the first lap of recording (instant fields) and fields that emerged on any subsequent lap (delayed fields).^15^ Early shifts were quantified as the mean MSD across the first 3 laps (Fig. S11) following field onset, and later shifting was quantified as the slope of MSD across laps 4 – 30.

### Principal components analysis

Principal components analysis (PCA) was used to characterize lap-by-lap differences in CA1 population activity in the familiar environment.^69^ PCA was performed on the mean activity of each cell across the first 15 laps of F1 and F2. Activity from individual laps was projected into this shared PCA space. Lap trajectory lengths were quantified as the total Euclidean distance between consecutive bins using the first 5 principal components (∼75% of population variance, Fig. S10). For individual mouse PCA, trajectories were smoothed using a 3-bin rolling average and trajectory lengths were normalized to the mean across all laps to account for differences in cell numbers (Fig. S10b-d). The effect of GC inhibition on trajectory length on lap 1 of F2 was evaluated using permutation testing (see Statistics).

### Population-vector template decoding

To decode episode identity and location from neural activity, we used a population vector-based template-matching decoding approach.^32,33,35^ The first 15 laps in each episode were used. For episode decoding, PV templates were created for each episode from the average activity across five randomly selected laps (Fig. S12), with new laps selected for each iteration. Population activity at each spatial bin of each test lap was correlated with the corresponding spatial bin of the F1 and F2 episode templates, and the episode with the higher correlation was assigned as the decoded episode. Decoder accuracy was calculated as the proportion of correctly classified bins. A correlation modulation index was also calculated as the median normalized difference between correct and incorrect episode template correlations, (correct – incorrect)/ (correct + incorrect), with larger values indicating stronger episode discrimination. Overall performance was evaluated across all F1 and F2 laps, while analysis was restricted to F1 or F2 laps to assess asymmetry in decoding errors. For location decoding,^33^ a 5-lap F1 template was compared to single lap activity from F1 (within-episode) or F2 (across-episode). For each true location bin, the template bin with the highest correlation was identified as the decoded location. Location decoding error was defined as the circular distance between the true and decoded location, and the mean location decoding error across bins was calculated for each lap. Decoder performance was averaged across 1,000 iterations with randomly selected template laps. For statistical testing, permutation testing used 1,000 shuffles, each containing 100 decoder iterations.

### Statistics

Statistical tests were calculated using MATLAB. Unless otherwise stated, tests were two-tailed with α = 0.05, and error bars and shaded regions represent median ± 95% confidence intervals. Violin plots show all individual data points, with kernel density estimates constrained to a valid range for each measure (i.e., -1 to 1 for correlation values) and kept consistent across comparable plots. LME models (see above) were used for single-cell analyses, and Wilcoxon signed-rank tests were used to evaluate the consistency of the effects of GC inhibition across mice. Lap-by-lap PV correlations, PCA, place field shifting, and PV decoding analyses were assessed using permutation testing with 1,000 condition shuffles in which cells were randomly assigned to the saline or DCZ condition. Two-sided p values were defined as the proportion of condition-shuffled differences with an absolute value that exceeded the observed effect.

## Acknowledgements

This study was funded by NIH F32-NS124752 (DG), and DP2-NS111657, R01-MH136274, RF1 NS127123, The Whitehall Foundation, The Searle Scholars Program, The Sloan Foundation, and University of Chicago startup funds (MS). We thank Reilly McClanahan and Jovangelis Gonzalez Del Toro for assistance with this project, Daniel Dombeck and Connor Davidson for feedback on this manuscript, Kimberly Christian for feedback on an earlier version of this manuscript, and the Tonegawa lab for providing the Dock10-Cre mice. Figs. 1a, e, and m were created using BioRender.

This manuscript is the result of funding in whole or in part by the National Institutes of Health (NIH). It is subject to the NIH Public Access Policy. Through acceptance of this federal funding, NIH has been given a right to make this manuscript publicly available in PubMed Central upon the Official Date of Publication, as defined by NIH. The content is solely the responsibility of the authors and does not necessarily represent the official views of the National Institutes of Health.

## Author contributions

Douglas GoodSmith: Conceptualization, investigation, formal analysis, visualization, writing -initial draft, writing – review & editing. Antoine Madar: Formal analysis. William H. Carson: Investigation. Mark E.J. Sheffield: Conceptualization, supervision, writing – review & editing.

