## Supplemental Figures for "The dentate gyrus controls hippocampal representational drift"

**Supplementary figures:**

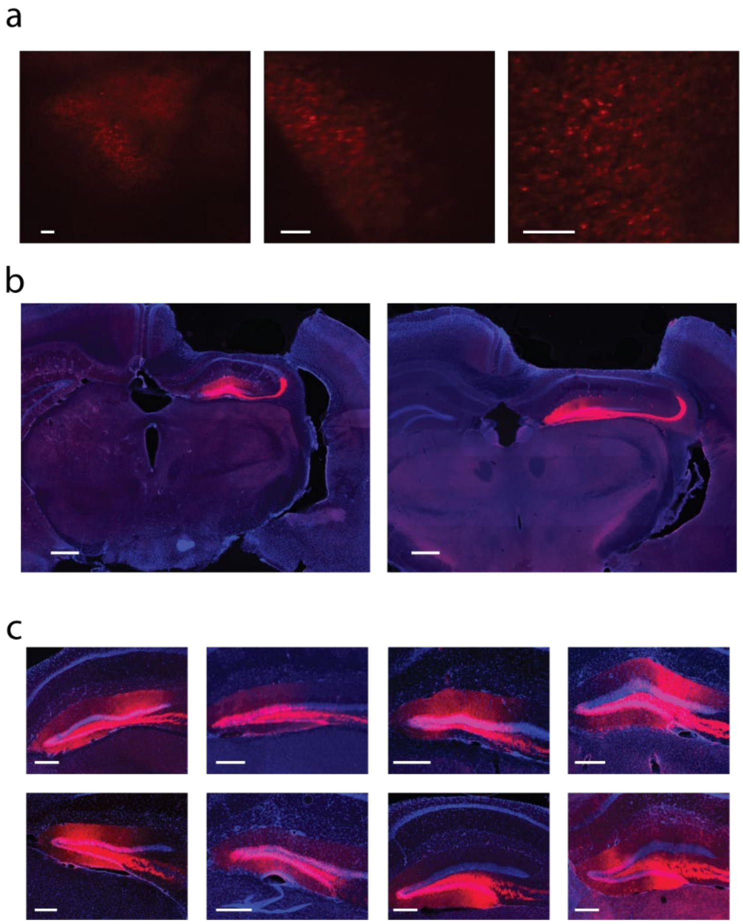


**Figure S1: Validation of inhibitory DREADD expression in granule cells**

(a) Sample FOVs from the granule cell layer imaged at 1064 nm confirming mCherry expression from the DREADD virus. Scale bars: 50μm.

(b-c) Representative histology sections showing mCherry expression (red) and DAPI staining (blue). mCherry expression was localized to the granule cell layer and mossy fiber projections to CA3. Scale bars: b, 500 μm; c, 250 μm.

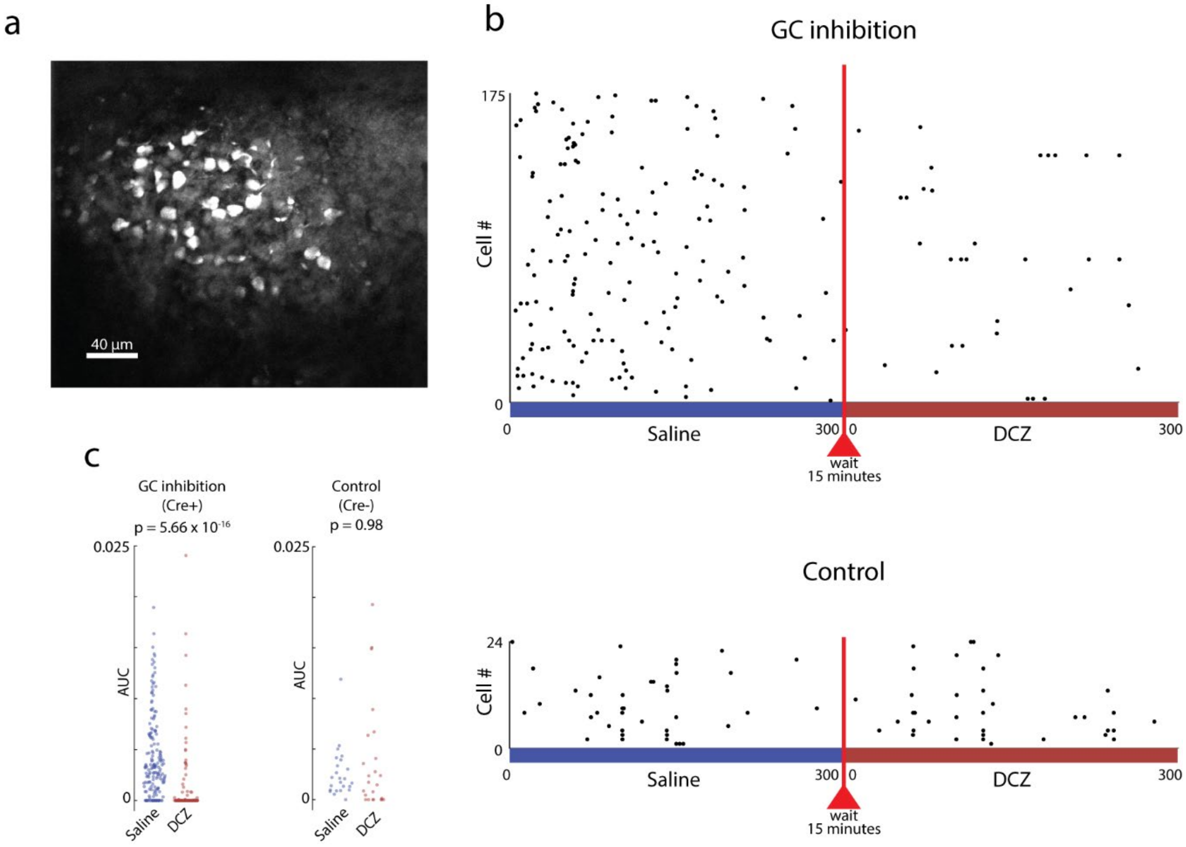


**Figure S2: DCZ injection significantly reduced granule cell activity**

DCZ was used over the more commonly used Clozapine N-Oxide (CNO) as DCZ is effective at activating DREADD receptors at much lower doses than CNO,[26,51] and due to the slow kinetics and known off-target effects of CNO.[52] Low doses of DCZ have been shown to produce significant DREADD-mediated effects within 10 minutes and persisting for at least 2 hours.[26] To confirm that the dose of DCZ used in this study was sufficient to inhibit granule cell activity, granule cell activity was directly recorded in two Dock10-Cre(+) mice and one Dock10-Cre(-) control mouse. All mice received the same virus injections described in Fig. 1a.

(a) Example GC imaging field of view (maximum projection).

(b) Raster plot showing calcium transient times for all 175 active granule cells recorded in GC inhibition mice (top) and all 24 active granule cells recorded in the control mouse (bottom). Mice were injected with saline 15 minutes before imaging. GC activity was recorded for ~5.5 minutes (blue) in the dark (no VR environment). Mice were then injected with 0.1 mg/kg DCZ while remaining head-fixed on the imaging rig (red line). GC activity was recorded again for ~5.5 minutes starting 15 minutes after DCZ injection (red).

(c) Area under the curve (AUC) of GC calcium transients. DCZ significantly reduced GC activity in GC inhibition mice (signed-rank test across cells, $p = 5.66 \times 10^{-16}$) but had no significant effect on the control mouse (signed-rank, $p = 0.98$).

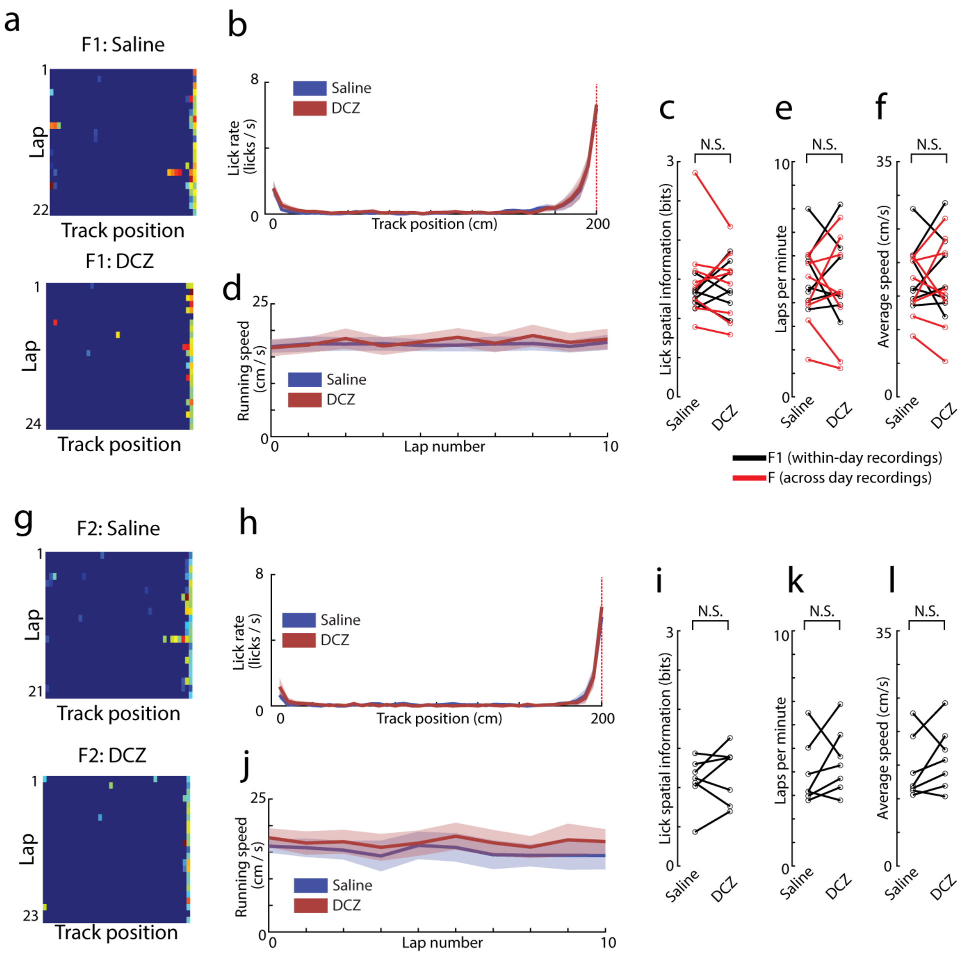


**Figure S3: GC inhibition did not affect behavior**

(a-f) Licking behavior and running speed in the first familiar episode of the day. Licking rate and running speed data from the within-day group (n = 7, F1 episode) and the across-day group (n = 8, full familiar episode) were pooled for analysis (n = 15 total episodes from 13 mice, data from two mice were included in both groups).

(a) Licking heat maps for one example mouse in the F1 episode on the saline (top) and GC inhibition (bottom, DCZ) days. Licking was concentrated near the end of the track, where reward was delivered, on both days.

(b) Mean licking rate across track positions, averaged across all laps. The mean and SEM across mice are shown for saline (blue) or DCZ (red) injected day. Licking is concentrated near reward delivery (dashed red line), with anticipatory licking immediately before reward delivery.

(c) The spatial information of licking (calculated as the mutual information of spatial bin and lick rate) was used as a measure of licking precision. GC inhibition did not significantly affect licking spatial information. Pooled across all mice, signed-rank $p = 0.98$; within-day recordings (black) signed-rank $p = 0.38$; across-day recordings (red) signed-rank $p = 0.38$.

(d) Mean running speed across the first 10 laps of the familiar episode. The mean and SEM across mice are shown for saline and DCZ days.

(e-f) GC inhibition did not affect running speed. Laps per minute (e) and average running speed (f) were not significantly different between saline and DCZ days (signed-rank tests; laps per minute: pooled $p = 0.97$; within-day $p = 0.98$; across-day $p = 0.97$. Running speed: pooled $p = 0.89$; within-day $p = 0.94$; across-day $p = 1$).

(g-l) Same as a-f, but for the F2 episode in the within-day recording group ($n = 7$).

(g-h) Licking was concentrated at the end of the track on both saline and DCZ days.

(i) GC inhibition did not affect licking spatial information (signed-rank $p = 0.47$).

(j) Mean running speed across the first 10 laps was consistent and similar between saline and DCZ days.

(k-l) GC inhibition did not affect laps per minute (k, signed-rank $p = 0.31$) or average running speed (l, signed-rank $p = 0.30$) in F2.

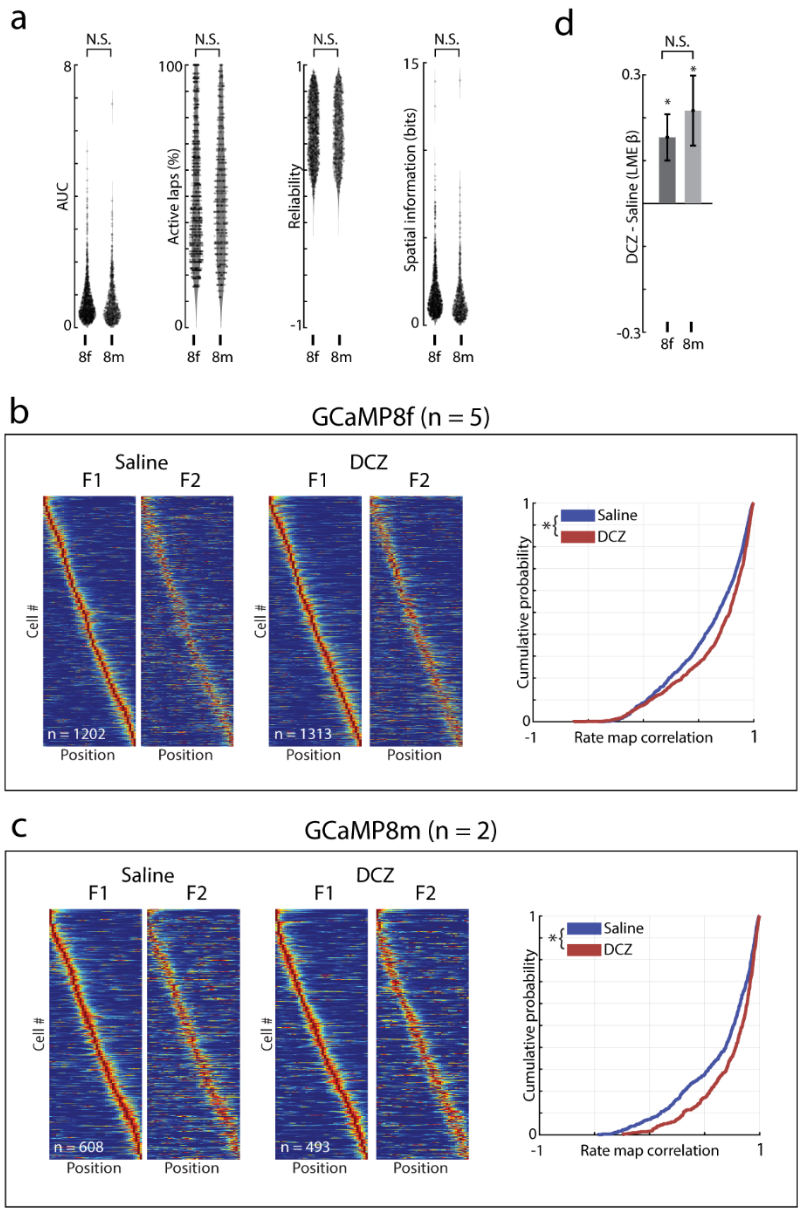


**Figure S4: Effects of GC inhibition were consistent across calcium indicators**

(a) Place cell activity measures were similar between mice imaged with GCaMP8f (n = 1,202 place cells from 5 mice) and GCaMP8m (n = 608 place cells from 2 mice). Activity properties were assessed in the F1 episode under the saline-injected condition. Significance was evaluated for each cell property with an LME including drug, calcium indicator, and drug x indicator interaction as fixed effects and mouse as a random effect. From left to right, there was no significant indicator effect on AUC ($\beta = 0.03$, $p = 0.76$), percent of laps with activity ($\beta$ = -0.006, p = 0.95), place cell reliability ($\beta$ = 0.04, p = 0.63), or spatial information score ($\beta$ = -0.21, p = 0.50).

(b) Effect of GC inhibition on CA1 population drift in GCaMP8f mice. Left, sorted place cell rate maps in the F1 episode and activity in the same cells in the F2 episode for saline and DCZ days. Right, GC inhibition significantly increased place cell rate map correlations between the F1 and F2 episodes relative to saline-injected control days (LME drug effect $\beta$ = 0.15, p = $2.18 \times 10^{-8}$).

(c) Same as b but for GCaMP8m mice. GC inhibition significantly increased place cell rate map correlations between F1 and F2 episodes (LME drug effect $\beta$ = 0.22, p = $2.15 \times 10^{-7}$).

(d) GC inhibition reduced CA1 place cell drift in both GCaMP8f and 8m mice, with no significant difference in the effect of GC inhibition between calcium indicators (drug x indicator interaction, $\beta$ = 0.06, p = 0.21).

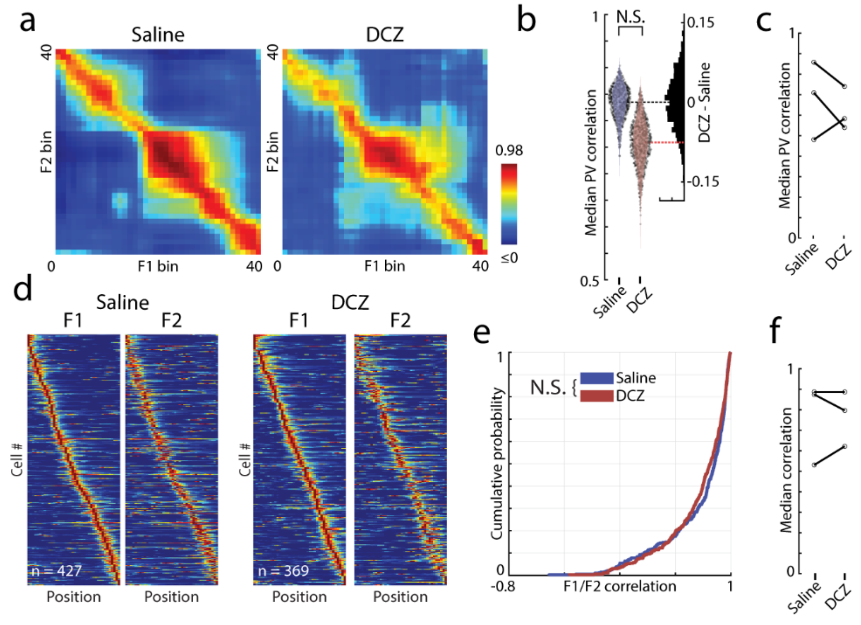


**Figure S5: DCZ injection did not affect drift of established representations in Cre-negative control mice.**

CA1 activity was recorded from three Cre-negative control mice. These mice received the same virus injections described in Fig. 1a and underwent the same imaging procedure (F1, N1, F2, N2 on separate saline and DCZ injection days) described in Fig. 1m.

(a) PV correlation matrices between F1 and F2 episodes following saline and DCZ injections.
(b) PV correlations between familiar episodes were not significantly affected by DCZ relative to saline (bootstrap permutation test $p = 0.17$).
(c) Median PV correlation values for each mouse.
(d) Sorted place field maps for saline and DCZ-injected imaging days.
(e) Place cell rate map correlations did not significantly differ between saline and DCZ-injected days (LME drug effect $\beta = -0.03$, $p = 0.48$).
(f) Median place cell correlation values for each mouse.

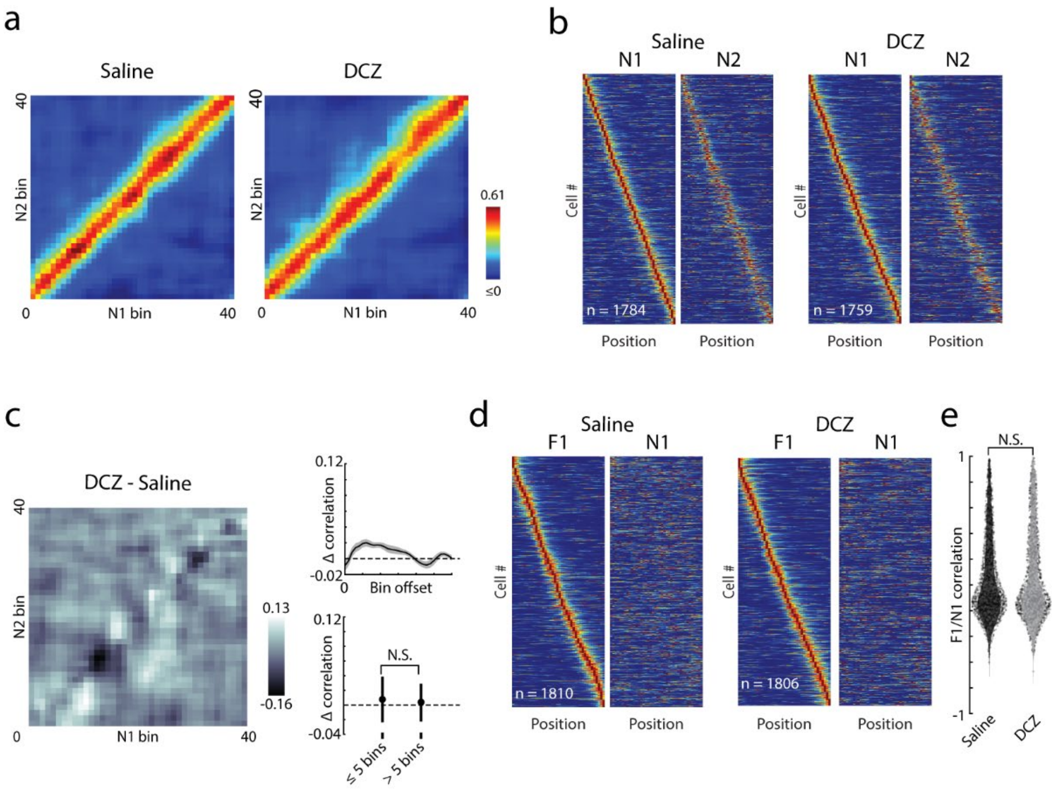


**Figure S6: GC inhibition did not affect stability of CA1 representations in the novel environment or remapping between environments**

(a-c) Effects of GC inhibition in the novel environment.

(a) PV correlation matrices between N1 and N2 episodes on saline and DCZ injection days. Saline: n = 6,375 cells; DCZ: n = 7,001 cells.

(b) Sorted place cell rate maps in N1 and N2 episodes on saline and DCZ injection days.

(c) Difference between PV correlation matrices for saline and DCZ conditions in the novel environment (left). PV correlation differences were not concentrated near the main diagonal (right). PV correlations for spatially similar bins (within 5 bins of main diagonal) were not significantly different from correlations for more distant bins (> 5 bins from the main diagonal, bootstrap permutation test, $p = 0.82$).

(d-e) Place field remapping between the familiar and novel environment.

(d) Place fields identified in the F1 episode are plotted by their peak activity in the F1 episode. The same cells are shown in the same order during the N1 episode, demonstrating significant reorganization of place cell activity (remapping) between environments.

(e) Place cell correlation values between F1 and N1 rate maps were not significantly affected by GC inhibition (LME drug effect $\beta = 0.02$, $p = 0.16$).

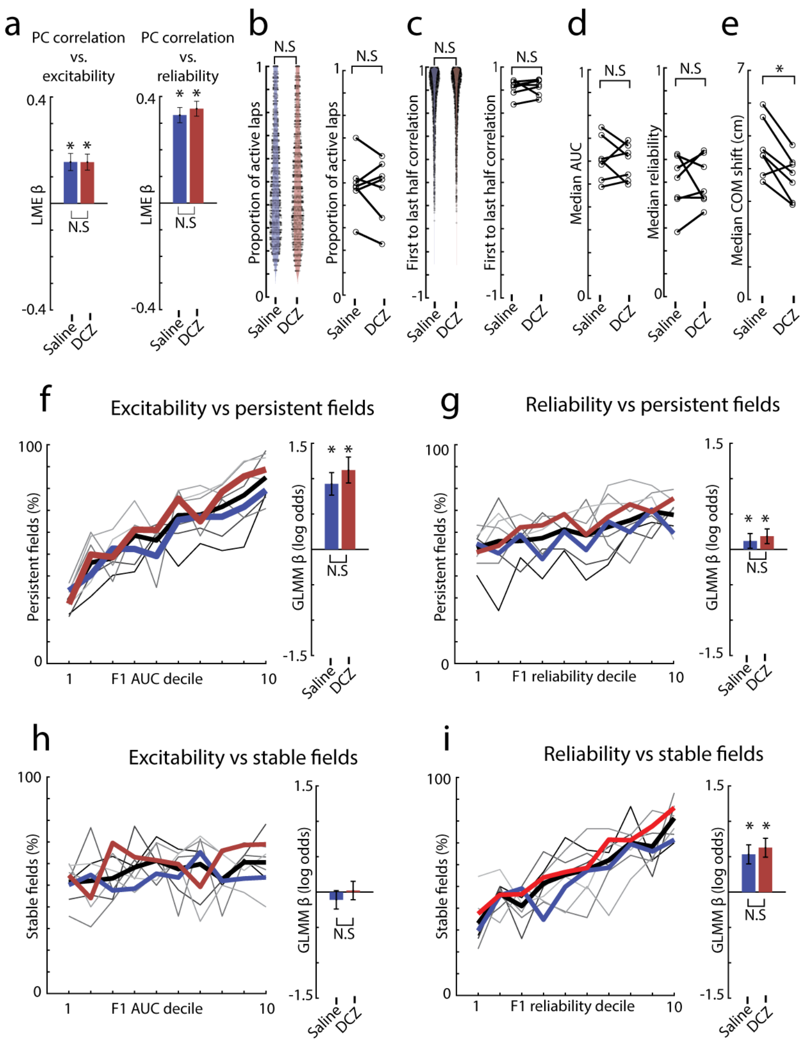


**Figure S7: Single-cell activity properties are associated with distinct components of representational drift.**

(a) Excitability (left, AUC) and reliability (right) were positively associated with place cell rate map correlations between familiar episodes in both saline and DCZ conditions, with no significant difference between conditions (Excitability: saline β = 0.16, p = 2.38 x $10^{-21}$; DCZ β = 0.16, p = 5.71 x $10^{-24}$; DCZ – saline β = -0.0004, p = 0.99. Reliability: saline β = 0.33, p = 2.25 x $10^{-106}$; DCZ β = 0.35, p = 2.77 x $10^{-125}$; DCZ - saline β = 0.02, p = 0.23).

(b) GC inhibition does not affect an alternate measure of place cell excitability, the proportion of laps with at least one calcium transient (LME β = 0.007, p = 0.82), or the per-mouse median of this measure (signed-rank p = 0.69).

(c) GC inhibition has no effect on an alternate measure of place cell reliability, the correlation between the average activity during the first and last halves of the F1 episode (LME β = 0.011, p = 0.63). The per-mouse median correlation value was also not affected by GC inhibition (signed-rank p = 0.30).

(d) Per-mouse median values for AUC and reliability were not significantly affected by GC inhibition (signed-rank tests: AUC, $p = 0.94$; reliability, $p = 0.69$).
(e) Median place field COM displacement was reduced by GC inhibition in 6/7 individual mice (signed-rank $p = 0.03$).
(f-i) Relationships between single-cell properties and components of drift. We defined place field persistence as the presence of a place field in F2 (at any location) among cells that had a detected F1 field. Among persistent fields, we defined spatial stability as fields with a COM displacement ≤1 bin. Excitability and reliability were binned into deciles, and the proportion of persistent or stable fields in each decile is plotted. Thin lines: individual mice; thick lines: pooled across mice (blue, saline; red, DCZ; black, pooled saline and DCZ).
(f) Place cell excitability was positively associated with field persistence under both saline and DCZ conditions, with no significant difference between conditions (saline $\beta = 0.93$, $p = 1.16 \times 10^{-29}$; DCZ $\beta = 1.12$, $p = 1.82 \times 10^{-32}$; DCZ – saline $\beta = 0.20$, $p = 0.11$).
(g) Place field reliability was positively associated with field persistence under both saline and DCZ conditions, with no difference between conditions (saline $\beta = 0.12$, $p = 0.03$; DCZ $\beta = 0.18$, $p = 4.64 \times 10^{-4}$; DCZ – saline $\beta = 0.07$, $p = 0.37$). Field persistence was more strongly associated with excitability than reliability (Fig. 2i).
(h) Place cell excitability was not significantly associated with the proportion of spatially stable fields under either saline or DCZ conditions, with no significant difference between conditions (saline $\beta = -0.11$, $p = 0.10$; DCZ $\beta = 0.02$, $p = 0.75$; DCZ – saline $\beta = 0.13$, $p = 0.17$).
(i) Place cell reliability was positively associated with the proportion of spatially stable fields under both saline and DCZ conditions, with no significant difference between conditions (saline $\beta = 0.53$, $p = 1.13 \times 10^{-14}$; DCZ $\beta = 0.63$, $p = 6.47 \times 10^{-20}$; DCZ – saline $\beta = 0.09$, $p = 0.33$). Field stability was more strongly associated with reliability than excitability (Fig. 2j).

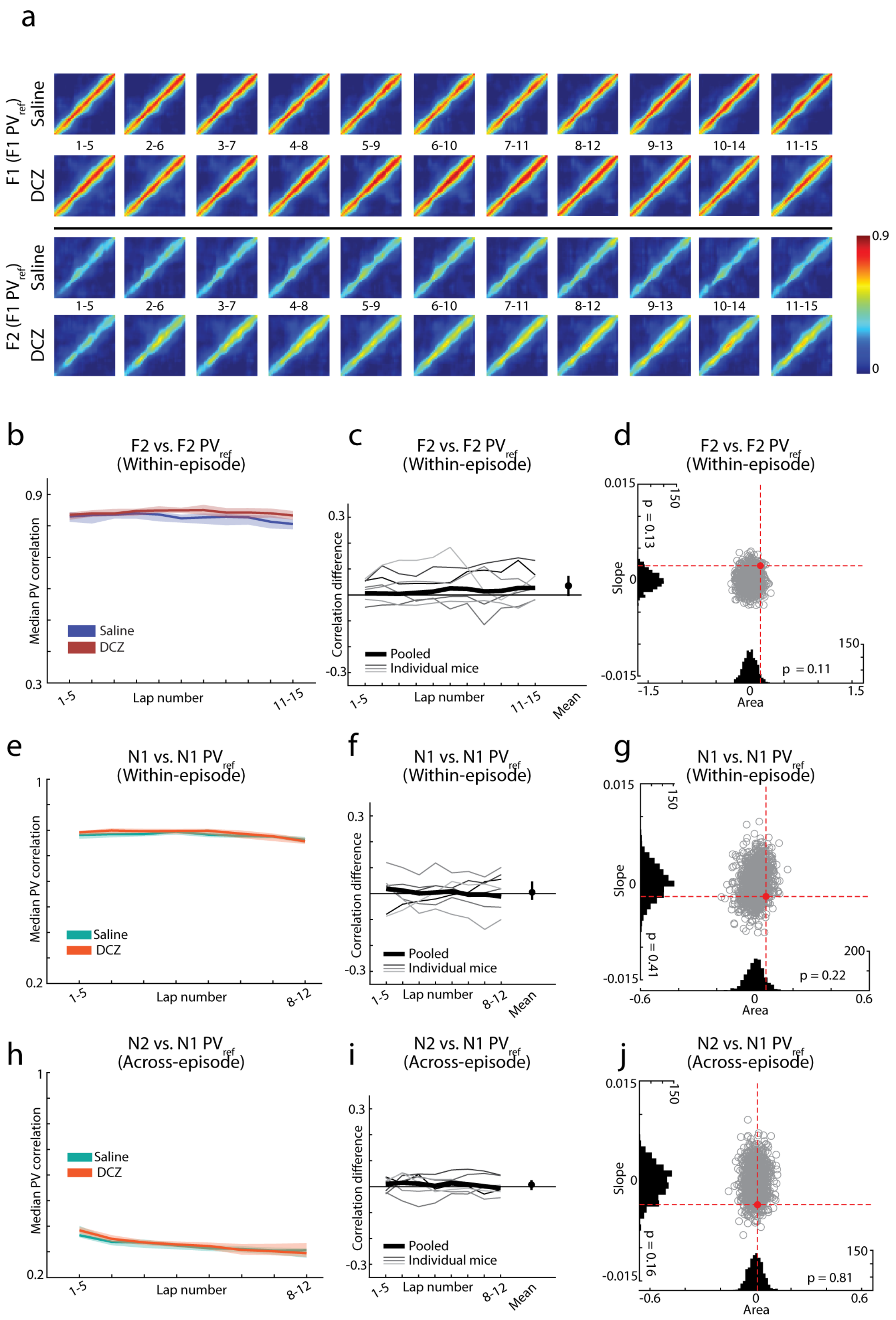


**Figure S8: Lap-by-lap PV correlations matrices and F2 within-episode PV correlations**

(a) PV correlations matrices between the F1 $PV_{ref}$ and rolling-average PVs from each lap of F1 (top) and F2 (bottom).

(b-d) Same as Fig. 3b-d, but comparing lap-by-lap activity in F2 to a $PV_{ref}$ created from the average population activity across the first 15 F2 laps. As in Fig. 3, within-episode PV correlations in F2 were not significantly affected by GC inhibition. The area between the saline and DCZ PV curves was not significantly different from the distribution of condition shuffled values (permutation test $p = 0.11$) and there was no significant slope in the difference between conditions to indicate an evolving effect throughout the episode (permutation test $p = 0.13$).

(e-j) Same as 3b-g, but for the novel environment. Comparing lap-by-lap activity in N1 (e-g) or N2 (h-j) to a $PV_{ref}$ created from the average population activity across the first 12 N1 laps. Within-episode (e-g) and across-episode (h-j) PV correlations in the novel environment were not significantly affected by GC inhibition. The area between the saline and DCZ curves and slope in the difference between conditions were not significantly different from a distribution of condition shuffled values (permutation tests, within-episode: area $p = 0.22$, slope $p = 0.41$; across-episode: area $p = 0.81$, slope $p = 0.16$).

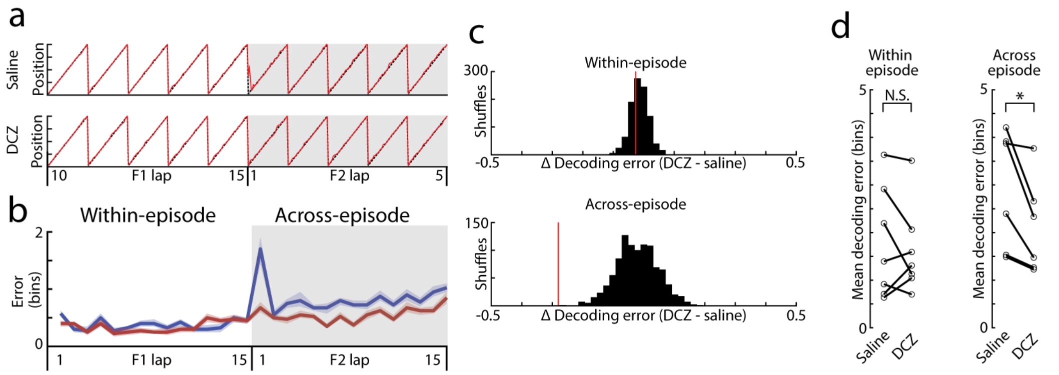


**Figure S9: PV template matching spatial location decoding.**

(a) Example decoded location (red line) and true location (dashed black line) for laps 10-15 of F1 and laps 1-5 of F2.

(b) Across 1,000 classifier iterations, template PVs were created from five random F1 laps, and location decoding error was calculated at each spatial bin for all laps not included in the template. Decoding error is shown within-episode (error in F1 laps) and across-episode (error in F2 laps) for both saline (blue) and DCZ (red). Mean and 95% confidence interval across classifier iterations.

(c) The average difference in decoding error between saline and DCZ conditions was not significantly different from a condition-shuffled distribution for within-episode decoding (top, permutation test $p = 0.41$), suggesting preserved within-episode spatial coding. Across-episode decoding errors (bottom) were significantly reduced following GC inhibition relative to the condition-shuffled distribution (permutation test $p < 0.001$), suggesting a greater similarity of F2 activity to the established F1 representation following GC inhibition.

(d) Decoders generated with data from individual mice revealed similar results, with no significant effect of GC inhibition on within-episode decoding (signed-rank $p = 0.69$) and significantly reduced across-episode decoding errors following GC inhibition across mice (signed-rank $p = 0.02$).

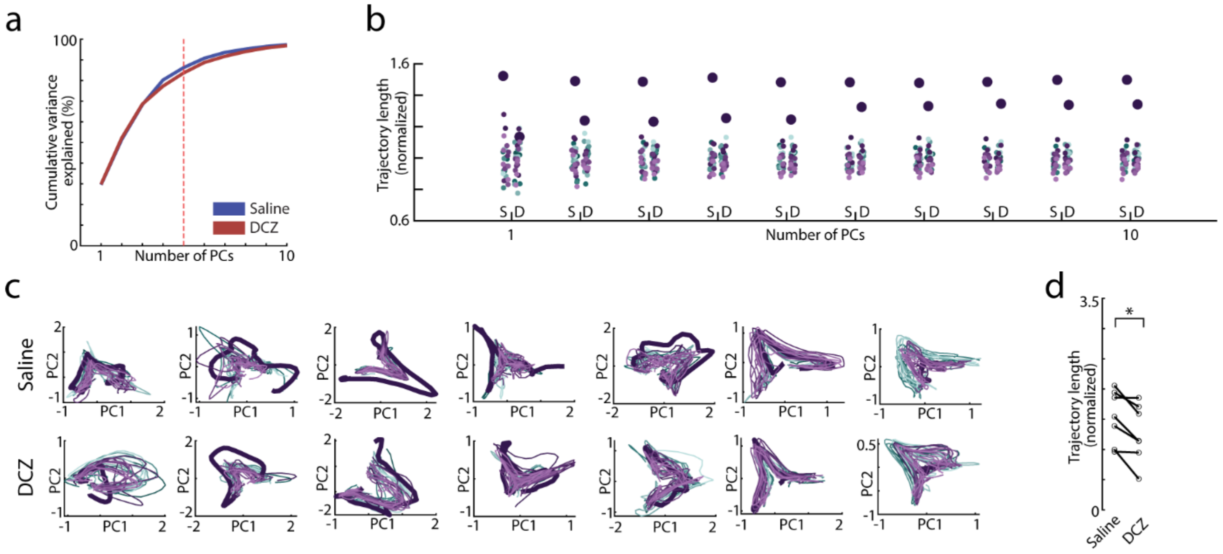


**Figure S10: PCA dimensionality and individual mouse PCA trajectories**

(a) Cumulative variance explained by PCA of population activity. The first 5 PCs (dashed red line) were used for trajectory analyses and accounted for roughly 75% of the total variance.

(b) The increased trajectory length on lap 1 of the F2 episode was consistently observed across PCA dimensionalities. Trajectory lengths for individual laps in the F1 (green dots) and F2 (purple dots) episodes are shown, measured using between 1 and 10 PCs. The large dark dot reflects the first lap of F2. For each number of PCs, trajectory lengths were normalized to the mean trajectory length across all laps to account for increasing trajectory lengths with dimensionality.

(c) PCA was performed separately on the population activity from each mouse on both the saline (top) and DCZ (bottom) days. The first 2 PCs are shown for each mouse, with F1 laps (green) and F2 laps (purple) shown. The thick purple line reflects the first lap of the F2 episode.

(d) F2 lap 1 trajectory length, normalized to the mean trajectory length of all laps, for individual mice. F2 lap 1 trajectory length was greater under the saline condition than the DCZ condition for all mice (signed-rank $p = 0.02$).

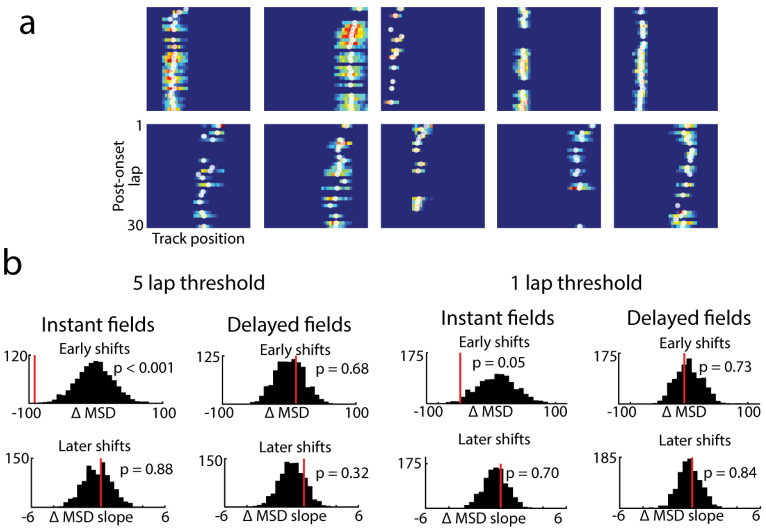


**Figure S11: place field shifting analysis**

(a) Example place field rate maps, showing the first 30 laps after field onset. Each lap's COM is shown as a white dot overlaid on the rate map.

(b) Reduced early shifting in instant fields following GC inhibition was not affected by the choice of laps to include in the early vs late shifting analysis. GC inhibition reduced the mean MSD of the first 5 laps (left, $p < 0.001$) and first lap (right, $p = 0.05$) but had no effect on the rate of later shifting or in delayed fields.

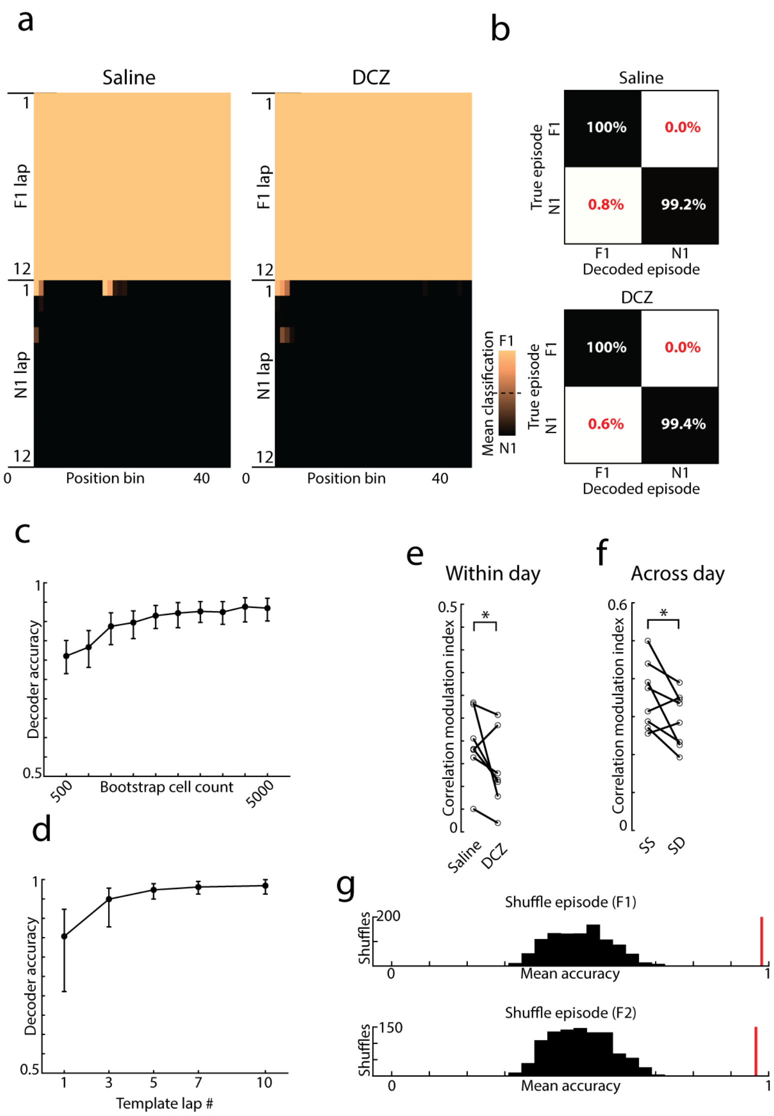


**Figure S12: Additional episode decoding analysis.**

(a-b) PV template-matching decoding between familiar (F1) and novel (N1) environments.

(a) Heatmaps showing the proportion of decoder iterations classified as F1 or N1 for each lap (row) and spatial bin (column). Lighter colors indicate more F1 classifications and darker colors indicate more N1 classifications. The first twelve laps of each episode were used for this analysis (the minimum number of laps across all mice in the novel environment).

(b) CA1 population activity could reliably decode between familiar and novel environments under both saline and DCZ conditions.

(c) Familiar episode decoder accuracy vs the number of cells used for decoding. Cells were bootstrap sampled with replacement to between 500 and 5,000 cells (in 500-cell increments), and decoder accuracy was calculated across 100 classifier iterations using randomly selected template laps. This procedure was repeated 1000 times. Mean decoder accuracy (percentage of correctly classified spatial bins) and 95% CI are shown. Decoder accuracy was significantly greater than chance with as few as 500 cells.

(d) Episode decoder accuracy vs the number of laps used to generate each template PV. Templates of 5 laps were used to maximize the tradeoff between accuracy and template size.

(e) Correlation modulation index for decoders created from individual mouse population activity. GC inhibition reduced decoder performance in 6 of 7 mice for the within-day (F1, N1, F2, N2) recordings (signed-rank $p = 0.047$).

(f) Same as e, but for cells tracked across days. GC inhibition reduced decoder performance for 6 of 8 mice (signed-rank $p = 0.039$).

(g) Decoder performance was compared to chance performance by randomly shuffling laps between F1 and F2 episodes and recalculating decoder accuracy (histograms). Despite reduced decoder performance following GC inhibition (Fig. 4), mean decoder accuracy was significantly above the shuffled distribution in both F1 (top) and F2 (bottom) episodes (permutation tests $p < 0.001$).
